# Chronic activation of astrocytic G_q_ GPCR signaling has causal effects on visual LTP formation: implications for neurodegenerative diseases

**DOI:** 10.1101/2024.04.16.589763

**Authors:** Elsie Moukarzel, Sharmilee Antoine, Sophie Guinoiseau, Bruna Rubino, Jacques Stinnakre, Cendra Agulhon

## Abstract

Astrocytes are the most abundant glial cells in the central nervous system and interact with other cell types, including neurons and microglia, *via* G_q_ protein-coupled receptors (G_q_ GPCRs) present on their surface. Astrocytic G_q_ GPCR activation induces Ca^2+^ release from internal stores, leading to intracellular Ca^2+^ elevations. There is emerging evidence supporting that astrocytic G_q_ GPCR Ca^2+^ elevations are upregulated and dysregulated in neurodegenerative diseases and are thought to play an important role in the pathogenesis of such diseases. Furthermore, astrocytic G_q_ GPCR Ca^2+^-dependent release of neuroactive or inflammatory molecules from astrocytes may occur in the early steps of the stress/inflammatory process in the diseased brain. In addition, low grade and chronic brain inflammation is involved in the etiology of neurodegenerative diseases.

We hypothesized that chronic activation of astrocytic G_q_ GPCR Ca^2+^ signaling leads to an altered production of glutamate or pro-inflammatory factors from astrocytes, and consequent deficits in synaptic transmission, long-term potentiation (LTP), and memory formation. To test this hypothesis, we used an AAV-based chemogenetic tool to selectively activate astrocyte G_q_ GPCR Ca^2+^ signaling combined with *in vivo* electrophysiology, immunohistochemistry, and biochemistry.

Using the mouse primary visual cortex (V1) as a model system, we found that chronically increased astrocytic G_q_ GPCR Ca^2+^ signaling leads to a decrease in LTP of visual-evoked potentials. Such LTP impairment was associated with microglial reactive phenotype - displaying a hyper-ramified and proliferative state - as well as a decrease in the number of interleukin 33 (IL-33)-expressing astrocytes. Our study is the first to have shown that chronic astrocytic G_q_ GPCR activation is sufficient to alter visual LTP and induce astrocyte-to-microglia communication, possibly through and IL-33 pathway in the adult brain. Because GPCRs are important drug targets, our study could have relevant therapeutic implications in the treatment of some neurodegenerative diseases.

## Introduction

Astrocytes are the major glial cell type in the central nervous system (CNS), interacting with nearly all neighboring cells, including neurons, microglial cells and blood vessels^1^. Until recently and for many years, they were considered as passive components sealing gaps between neurons, and providing them metabolic support^2^. However, the use of modern tools has helped investigators to unravel novel crucial astrocytic roles in synapse formation, maturation, and elimination as well as in neuronal activity^3–5^.

Astrocytes are continuously aware of their surrounding environment and express an array of G protein-coupled receptors (GPCRs) at their plasma membrane^6–10^, allowing them to sense neuroactive or pro-inflammatory molecules released by neurons and other cell types^11^. Astrocytic G_q_ GPCR activation induces intracellular Ca^2+^ elevations *via* inositol 1,4,5-trisphosphate receptor (IP_3_R)-mediated Ca^2+^ efflux from the endoplasmic reticulum^12^. Furthermore, it has been shown that under certain experimental conditions, astrocytic G_q_ GPCR Ca^2+^ elevations lead to the release of neuroactive molecules (called gliotransmitters), such as ATP^13,14^, glutamate^15–17^, D-serine^18^, and GABA^19–21^, modulating synaptic transmission, plasticity, memory and complex behavior^22^. Together these data led to the concept of gliotransmission – defined as the rapid and regulated release of gliotransmitters by astrocytes in an activity and G_q_ GPCR, Ca^2+^-dependent manner. Although the physiological relevance of this mechanism is under debate^23–26^, there is emerging evidence supporting Ca^2+^-dependent gliotransmitter release in the early steps of the stress/inflammatory process in the diseased or damaged brain^26,27^. For instance, inflammatory mediators are sufficient to induce the release of glutamate and pro-inflammatory mediators from astrocytes through activation of G_q_ GPCRs^28,29^. Additionally, upregulated and dysfunctional astrocytic G_q_ GPCR (purinergic P2Y_1_R and glutamatergic mGluR5)-mediated Ca^2+^ signaling are thought to play an important role in the onset or pathogenesis of Alzheimer’s disease (AD)^30–34^. Moreover, there is accumulating evidence that low grade and long-lasting brain inflammation is involved in the etiology of AD^35–38^. Finally, astrocytes play critical roles in the progression and maintenance of brain inflammation by producing and responding to stress/inflammatory mediators^39^.

We hypothesized that during adult life challenges, chronic stress/inflammation-induced astrocytic G_q_ GPCR Ca^2+^ activation may trigger upregulated production of glutamate and/or pro-inflammatory molecules in astrocytes. All of these could consequently cause long-term alterations in excitatory synaptic transmission and plasticity, and thus contribute to the early sequences of events contributing to hyperexcitability, excitotoxicity, synapse loss, neurodegeneration, and thus to the pathogenesis of AD.

The main goal of this study was to investigate whether *in vivo* chronic astrocytic G_q_ GPCR activation was sufficient to induce AD-like long-term potentiation (LTP) and memory impairments. The mouse primary visual cortex (V1) was chosen as a model system, a cortical region in which astrocyte function^40^ as well as visual processing have been found to be impaired in neurodegenerative diseases such as AD^41–51^.

To address this question we selected the DREADD (designer receptors selectively activated by designer drugs) technology^52,53^ for which we pioneered the use in astrocytes to specifically activate G_q_ GPCR signaling cascades in these cells^22^. Such technology was combined with adeno-associated viral (AAV) tools to target astrocytes selectively in V1 of adult mice. State-of-the-art *in vivo* electrophysiology, biochemistry, and immunohistochemistry were then used to measure neuronal, astrocytic and microglial activity changes.

We found that chronic (several week-long) and selective astrocytic G_q_ GPCR activation in mouse V1 induces a decrease in LTP of visual-evoked potential (VEPs). Such LTP (also called stimulus-selective-response potentiation or SRP) is known to normally occur upon repeated visual grating stimulus presentations, and to reflect a form of visual perceptual memory exclusively stored in V1^54,55^. The astrocytic G_q_ GPCR-induced LTP deficit here described was not associated with any changes in the number of thalamocortical excitatory synapses in V1 layer IV, but was accompanied by microglial cell reactivity, proliferation, and hyper-ramification as well as a decreased in the number of interleukin 33 (IL-33)-expressing astrocytes. Together these data suggest that visual cognition deficit found in AD^45–49^ might be due to chronic and upregulated astrocytic G_q_ GPCR activation^30–33^, possibly involving an IL-33-mediated astrocyte-to-microglial communication^56–59^.

To the best of our knowledge, our study is the first to have shown that chronic astrocytic G_q_ GPCR activation is sufficient to (i) alter visual LTP, (ii) mediate astrocyte-to-microglia communication possibly through and IL-33 pathway in the adult brain. These findings could have profound implication in understanding the role of astrocytes during early visual impairments in neurodegenerative disease, such as AD

## Material and methods

### Animals

Animal care and procedures were carried out according to the guidelines set out in the European Community Council Directive. The protocol was approved by the Committee on the Ethics of Animal Experiments of Paris Descartes University (Protocol number: 2020061613287484). Both male and female mice were group housed (5 mice/cage) on a 12h dark/light cycle at a temperature of 20 ± 1°C, with water and food provided *ad libitum*, humidity ∼50%. IP_3_R2-KO mice^60^ were obtained from J. Chen (University of California at San Diego). Adult C57BL/6JRj mice were from Janvier Labs. Furthermore, CAG-lox-STOP-lox-GCaMP6f transgenic mouse line was crossed with GLAST-CreERT² mouse^61^, to obtain a double transgenic mouse line GLAST-CreERT^2^::GCaMP6f^62^ used for Ca²^+^ imaging. All experiments were conducted during the dark phase. For immunohistochemistry and western blots, data were collected from 8–15-week-old mice, and from 14–15-week-old mice for *in vivo* recordings. All experiments were performed double blindly without knowledge of their genotype, or treatments.

### Stereotaxic AAV-or ACSF-injections

6-week-old male and females were anesthetized by intraperitoneal injections of ketamine/xylazine (100mg/kg and 10mg/kg) and placed in a stereotaxic apparatus (Kopf Instruments). After hair trimming, and before any surgical incision, eye-ointment (Ocrygel®) was applied generously on the animal eyes, and 1% lidocaine was applied over the head. 10% Povidone-iodine solution was used to clean the scalp. Then, 1cm antero-posterior long incision was made, followed by craniotomies using a 30-gauge needle without thinning the skull as it can easily be removed.

Injections were made in V1 using a nano-injector (Drummond ®), guiding a glass pipette filled with 0.5µL of solution. The viral solution was delivered at a rate of 0.05µL/min, during 10min (217 pulses, 2.3nL/pulses). Glass pipettes were left in place for at least 10 min to prevent virus reflux. Incision was sutured with 4-0 silk sutures (Ethicon ®). Animal temperature was monitored throughout the whole surgery, using a heating pad, and maintained at 37°C. For immunohistochemistry experiments, unilateral injections of AAV-8-GFAP-G_q_ DREADD were made at +/- 3.0 mm lateral to lambda and 400µm from the brain surface. For Western Blot and Ca²^+^ imaging experiments, two unilateral injections (to ensure a correct virus spread) of AAV-8-GFAP-G_q_ DREADD were made at +/- 2.7 and +/-3.3 mm lateral to lambda and 400µm from the brain surface. For *in vivo* electrophysiological recordings, two bilateral double of AAV-8-GFAP-G_q_ DREADD or artificial cerebrospinal fluid (ACSF) solution, were made at +/- 2,7 and +/-3,3 mm lateral to lambda and 400µm from the brain surface. Following surgery animals recovered overnight in low-heated cages.

### Electrode implant surgery

14-week-old male and females were anesthetized by intraperitoneal injections of ketamine/xylazine (100mg/kg and 10mg/kg) and placed in a stereotaxic apparatus (Kopf Instruments). After hair trimming, and before any surgical incision, eye-ointment (Ocrygel®) was applied generously on the animal eyes, and 1% lidocaine was applied over the head. 10% Povidone-iodine solution was used to clean the scalp. Then, 3 cm antero-posterior long incision was made. Skull was cleaned with hydrogen peroxide and periosteum was removed. A steel headpost was attached to the skull anterior to bregma using cyanoacrylate glue. Then, using a 26G needle, 2 holes (<300µm) were made in the skull (+/-2mm lateral; -0.75mm posterior to bregma) and custom-made silver wire reference electrodes were placed. Two other holes were made using a 30G-needle over binocular V1 (+/- 3.0 mm lateral to lambda) and 75µm diameter-wide tungsten recording electrodes (FHC), dipped in micro-fluorescent red spheres (FluoSpheres™ Carboxylate-Modified Microspheres, 0.1 µm, red fluorescent (580/605), 2% solids; Invitrogen #F8801) were implanted within each hemisphere, at 450µm depth from cortical surface. Red fluorescent beads allow to check the electrode track at the end of the experiments (**Supplementary Figure 2**). All implants were then secured in place using only cyanoacrylate glue to form a glue platform, and Zip Kicker® was applied to ensure a stable cap. Sutures in the back and at the front were made. Mice were monitored postoperatively for signs of discomfort and were allowed 24 h for recovery.

### Preparation for brain slices and Ca²^+^ imaging

Briefly, GLAST-CreERT^2^::GCaMP6f brains injected with AAV-8-GFAP-G_q_ DREADD two weeks prior experiment, were extracted, and 400µm thickness slices containing V1 were made using a vibratome (Leica), at a 0.05 mm/s speed, 0.75mm amplitude, and 60 Hz vibrating frequency. Slices were then incubated 20min in 35°C ACSF containing (in mM) : 124 NaCl, 3 KCl, 1.25 NaH_2_PO_4_, 26 NaHCO_3_, 20 dextrose, 1 MgCl_2,_ 2 CaCl_2_. Then slices were transferred at room temperature for 45min, and then placed in the recording chamber of a custom-built 2-photon laser-scanning microscope with a 20x water immersion objective (x20/0.95w XLMPlanFluor, Olympus). GCaMP6f was excited at 920 nm with a Ti:Sapphire laser (Mai Tai HP; Spectra-Physics Newport Mountain View, CA). Brain slices were continuously superfused at a rate of 4 ml/min with ACSF and bubbled with 95% O2–5% CO2. To evoke intracellular Ca^2+^ elevations in GCaMP6-G_q_ DREADD-expressing astrocytes, CNO (10µM) was bath applied for 2 min.

### Western Blot

Animals were sacrificed one day after the end of the treatment (CNO 1mg/kg or Saline 0.9%) and brain were extracted, snap frozen at -30°C and stored at -80°C. Primary visual cortices from both hemispheres were punched (2x1mm diameter) and homogenized in 150µL RIPA buffer (50mM Tris pH 8, 150mM NaCl, 1% NP-40, 0.5% deoxycholate, 0.1% SDS, 1 mM sodium orthovanadate) with 1X proteases inhibitor cocktail COmplete (Roche) and 1X Halt phosphatase inhibitor cocktail (Thermo Scientific). Then, tissue lysate was extracted using a Bioruptor sonication system (Diagenode) and 2000 RPM centrifugation for 5 min, and only supernatants were kept. Protein concentration was assessed using a BCA assay kit (Bio-Rad) and equal amount of protein (25µg) from each sample was deposited and run on a 10% acrylamide SDS-page to separate proteins. Proteins were then transferred onto nitrocellulose membranes. Depending on protein weight, membranes were cut into horizontal strips to maximize the number of antibodies that could be assessed on each blot. Blots were saturated in 5% non-fat milk in TBS-Tween for 30min at room temperature under slow agitation. Membranes are then incubated with primary antibodies at 4°C overnight (**Supplementary Table 1**). The following day, membranes were washed three times in TBS-T, and incubated with corresponding peroxidase-labeled secondary antibodies (**Supplementary Table 1**) diluted in TBS-T for 1h30 with agitation. Bands were developed and visualized using Clarity ECL chemiluminescence technology (Biorad) and ImageQuant LAS4000 (GE Healthcare Life Sciences). On average, membranes were exposed between 30s and 2min, and the image just before saturation was used for quantification. For reprobing with different antibodies (in our case for tubulin), blots were washed three time in TBST, stripped with a buffer solution (0.5M Tris-HCl pH 6,7; 10% SDS, β−mercaptoethanol, deionized water) at 50°C for 30min, washed three times in TBS-T and then blocked again in 5% milk/TBS-T prior to re-incubation in anti-tubulin primary antibody overnight before following the same protocol as above. Each mean grey value corresponding to the signal was extracted using ImageJ software (NIH). Western blot experiments were replicated 4 times in order to dampen technique-dependent variability. In each replicate, the order of the deposits was changed to avoid any position effect within gels. For normalization, each value from the control (DREADD/Saline) and the experimental (DREADD/CNO) group was normalized to its corresponding tubulin mean grey value. Then, the 4 mean grey values obtained from the 4 replicates for the same sample were averaged. The obtained values were normalized to the corresponding non-injected hemisphere values, and finally normalized to the average value from the control group (DREADD/Saline). Moreover, to check that the punched region in the injected hemisphere expresses the hM3D_q_ receptor, WB was also performed to detect mCitrine expression levels. Only samples showing an mCitrine expression were kept for the analysis. Western Blot analyses were performed double-blind.

### Immunohistochemistry

Animals were transcardially perfused with 4% paraformaldehyde dissolved in 0.02 M phosphate buffer saline (PBS) under ketamine/xylazine (100mg/kg and 10mg/kg), anesthesia. Brains were removed, post-fixed for 24 h in 4% paraformaldehyde. Then, tissues were cryoprotected overnight at 4°C in 0.02 M PBS (pH 7.4) containing 20% sucrose, and frozen at -30°C in optimal cutting temperature compound. Sixteen μm thick sections were cut using a cryostat (Leica), mounted on Superfrost glass slides and stored at -80°C. The day of the experiment, sections were washed 3 times for 15 min each in 0.02 M PBS. For IL-33 staining, antigen retrieval was performed by incubating slides in antigen retrieval buffer (10 mM sodium citrate, 0.05% Tween-20, pH 6.0) at 96°C during 2min in a micro-wave and then 10min in the hot buffer. All sections were then incubated overnight in 0.02 M PBS containing 0.3% Triton X100, 0.02% sodium azide and primary antibodies (**Supplementary Table 1**) at room temperature in a humid chamber. To better visualize fine subcellular compartments expressing cytosolic mCitrine protein and to not miss any transgene expression, mCitrine and GCaMP6f signal were amplified using an antibody directed against GFP (1:1000, Invitrogen A10262. The following day, sections were washed 3 times for 15 min each in 0.02 M PBS and incubated for 2h at room temperature with corresponding secondary antibodies diluted in 0.02 M PBS containing 0.3% Triton X100 and 0.02% sodium azide (**Supplementary Table 1**). Then, sections were finally washed 3 times for 15 min in 0.02 M PBS and mounted between slide and coverslip using Vectashield medium containing DAPI (Vector Laboratories) or without DAPI for synapse counting experiment. Negative controls*, i.e*. slices incubated with secondary antibodies only, were used to set criteria (gain, exposure time) for image acquisition in each experiment. For VEPs experiments, all DREADD mice were sacrificed at the end of the experiment to check if the virus was expressed and if the electrode was correctly implanted within the viral spread (**Supplementary** Figure 2).

### Image acquisition, cell counting and mean density measurements

Image acquisition was performed with an Axio Observer Z1 epifluorescence Zeiss microscope, an ORCA Flash 2.8-million-pixel camera, and a PlanNeoFluar 20x/0.5NA objective. Two-to-four coronal sections per mouse were taken for analysis. Images were extracted using the ZEN 2011 blue edition software (Zeiss). Cell counting measurements were performed using manual cell counting plugin function in ImageJ software (National Institutes of Health, USA). To characterize the AAV-8-GFAP-G_q_ DREADD, cells were counted on 5-10 mice, 2 to 4 brain slices/mouse. For mean grey value measurement and cell counting, 2 to 4 slices were imaged for both hemispheres. To reduce mouse variability, each value obtained from each injected hemisphere was normalized to the value obtained from the non-injected hemisphere. To define the region to analyze, a region of interest (ROI) was chosen where the G_q_ DREADD was expressed (for the injected hemisphere), and a similar control area for the non-injected hemisphere. After normalizing to the non-injected hemisphere, values obtained from different slices for each mouse were averaged and then normalized to the control group average. For IL-33 staining across cortical layers, binned mean grey values at 50µm-depth intervals were extracted from one slice per mouse through ImageJ software using a line scan plugin. Data were extracted from brain surface, until the end of the viral spread. Values were averaged, normalized to the non-injected hemisphere, and plotted on a graph. To count the number of IL-33-positive cells, we counted the number of IL-33-positive nucleus. Therefore, only cells expressing IL-33 within the nucleus were taken into account.

### Sholl analysis

Sholl analysis was used to assess microglia complexity through their branching. 10 cells per hemisphere (non-injected and injected) from 2 slices were randomly selected. The line segment tool was used to draw a line from the center of each soma to the tip of its longest process, providing the maximum process length. Then the Concentric circles plugin on ImageJ was used with the first shell set at 2µm and subsequent 6 shells set at 8µm step sizes, to determine manually the number of intersections at each Sholl radius. Only microglial cells displaying a cell body and branches in the focal plan were chosen for analysis. Averaged values per hemisphere for each radius (DREADD/Saline and DREADD/CNO injected and non-injected hemispheres) were graphed, and the total number of intersections, normalized to the non-injected hemisphere and to the mean of the control group was also graphed. Soma size was also measured, and the number of microglial cells was counted to determine the average density.

### Synapse counting

To count the number of excitatory synapses, 2 to 4 coronal slices per mouse containing mCitrine staining (*i.e* AAV-8-GFAP-G_q_DREADD) and their respective contralateral control slices in V1 were imaged for the analysis, using a 63x-oil objective on a Zeiss LSM710 confocal microscope. 4.63 µm thickness z-stack images (optical section depth 0.63 μm, 8 slices/z-stacks from the surface, 1942 x 1942 pixel/image/scan) were taken. Exposure was set using the negative control slice. Images were deconvoluated using AutoQuant X3 software to remove background and improve resolution. To count the number of VGlut2 and Homer1 and colocalized puncta, IMARIS 9.6 software (Bitplane) was used. A machine learning program within the spot detection function was setup to automatically detect VGlut2 and Homer1 puncta with a diameter >0.2µm. Using a colocalization spot plugin, we defined a maximum center-to-center distance of 0.45µm between VGlut2 and Homer1 puncta to consider a synapse.

### *In vivo* chronic CNO treatment

CNO was dissolved in 0.01% DMSO and 0.9% saline to obtain a 1mg/kg/day dose of CNO. 2 weeks after stereotaxic AAV-8-GFAP-G_q_ DREADD or ACSF injections, CNO (1mg/kg) or saline (0.9%) was subcutaneously administrated once a day for 6 weeks to DREADD mice.

### Tamoxifen treatment

GLAST-CreERT^2^::GCaMP6f mice were intraperitoneally injected with tamoxifen (1 mg/day, Sigma) diluted in corn oil (Sigma) during 3 consecutive days. Animals were then used 2 weeks after the first day of treatment. All lines used were kept heterozygous for transgenes encoding GCaMP6f, Cre or CreERT^2^. Experiments were conducted in 6 weeks to 14-week-old male and female mice from the C57BL/6JRj background.

### Visual stimulus delivery

Visual stimuli were generated using Vision Work software (Vision Work Graphics, USA) to control stimulus drawing and timing. Visual stimuli was projected on a cathode ray tube television monitor situated 20 cm in front of head-fixed mice, therefore occupying 92° × 66° of the visual field. Mean screen luminance was 27 cd/m2. Visual stimuli consisted of full-field sinusoidal grating stimuli phase reversing at a frequency of 1 Hz (0.05 cyc/degree, 100% contrast). Grating stimuli covered the full range of monitor display values between black and white, with a y correction to guarantee constant total luminance in both grey screen and grating stimulus settings. Experiments were completely computerized, and visual stimulus consists of 5 min grey screen followed by 5 blocks of stimulation. Each stimulus block consisted of 200 phase reversals interleaved with 30s grey-screen of similar luminance. Before recording, mice were habituated to the head-fixed position and screen brightness by sitting in front of a grey screen for 30 min on each of 2 consecutive days. During the other 9 days of stimulation, 45° oriented visual stimulation were presented (familiar stimulus). On the 10^th^ day of stimulation, a new stimulus was presented (new stimulus 135°).

### *In vivo* electrophysiology data acquisition and analysis

Visual-evoked potential (VEP) recordings were conducted in awake, head-fixed mice. Two weeks before electrode implant surgery, mice were habituated to the electrophysiological setup. They were handled and explored the environment 10 min per day. Two recording channels were dedicated to VEPs recordings from V1 for each implanted hemisphere. All data were extracted and analyzed using Spike2 software (Cambridge electronic designed limited, CED). VEPs were averaged throughout all phase reversals within each block and peak-to-peak difference measured during a 500-ms period from phase reversal.

For SRP assessment in DREADD mice, only mice with the electrode implanted in the middle of a large viral spread were kept (**Supplementary** Figure 2).

### Statistical analysis

Data were acquired and analyzed blind of genotype and treatment. Data are shown as mean ± S.E.M. n values are either hemispheres, slices or mice, as indicated in each figure.

All statistical tests were performed after verifying data normal distribution using Shapiro-Wilk normality test. If normality assumptions were not met, non-parametric tests (two-tailed Mann-Whitney for 2 group analysis) were performed. If normality was met, parametric tests (two-tailed Student t-test for 2 group analysis) were used. For in vivo electrophysiology experiments, and plot scan analysis, Mixed-effects model (REML) followed by Tukey post-hoc test and Bonferroni test were used, respectively, because of some missing data points. For Sholl analysis experiment, 2-way ANOVA, was done followed by Bonferroni post-test w. All statistical tests were performed using GraphPad 8®. No statistical methods were used to pre-determine sample sizes but there are in the same range to those generally used in the field.

## Results

### AAV8-GFAP-G_q_ DREADD allows selectively DREADD expression in V1 astrocytes

To investigate the role of G_q_ GPCR Ca²^+^ signaling we used the G_q_-coupled designer receptor hM3D_q_ (G_q_ DREADD). To ensure regional and cellular specificity, we stereotaxically delivered in V1 layer IV, a serotype 8 AAV (AAV8) vector, encoding a Human influenza hemagglutinin epitope tagged protein (HA) fused with hM3D_q_ and a cytosolic mCitrine protein, under the control of the astrocytic glial fibrillary acidic protein (GFAP) promoter (AAV8-GFAP-G_q_DREADD) (**Figure 1A**). Two weeks after AAV injection, brain were extracted and immunohistochemistry was performed. mCitrine expression pattern (reflecting G_q_ DREADD-expressing cells) showed that G_q_ DREADD expression was limited to V1, and present in almost all V1 layers (**Figure 1B,C**, **Supplementary Table 2**, spread distribution: dorsoventral average = 614µm; mediolateral average = 746µm, anteroposterior average = 528µm; n=13 mice). Moreover, G_q_ DREADD expression was specifically found in 71% of S100β-expressing astrocytes (**Figure 1D,E**, **Supplementary Table 2**, n=10 mice), and was almost completely specific to astrocytes (>97% G_q_ DREADD-expressing cells were also expressing S100β; n=10 mice, **Supplementary Table 2**). Co-staining with NeuN neuronal marker (**Figure 1F**, n=10 mice), Iba1 microglial marker (**Figure 1G**, **Supplementary Table 2,** n=10 mice), and NG2 oligodendrocyte precursor marker (**Figure 1H**, **Supplementary Table 2**, n=5 mice) showed no overlap with G_q_ DREADD expression, thus confirming the astrocyte G_q_ DREADD specific expression. HA immunostaining revealed that G_q_ DREADD was expressed in the very fine processes of mCitrine-expressing astrocyte (**Figure 1E**).

**Figure 1:**
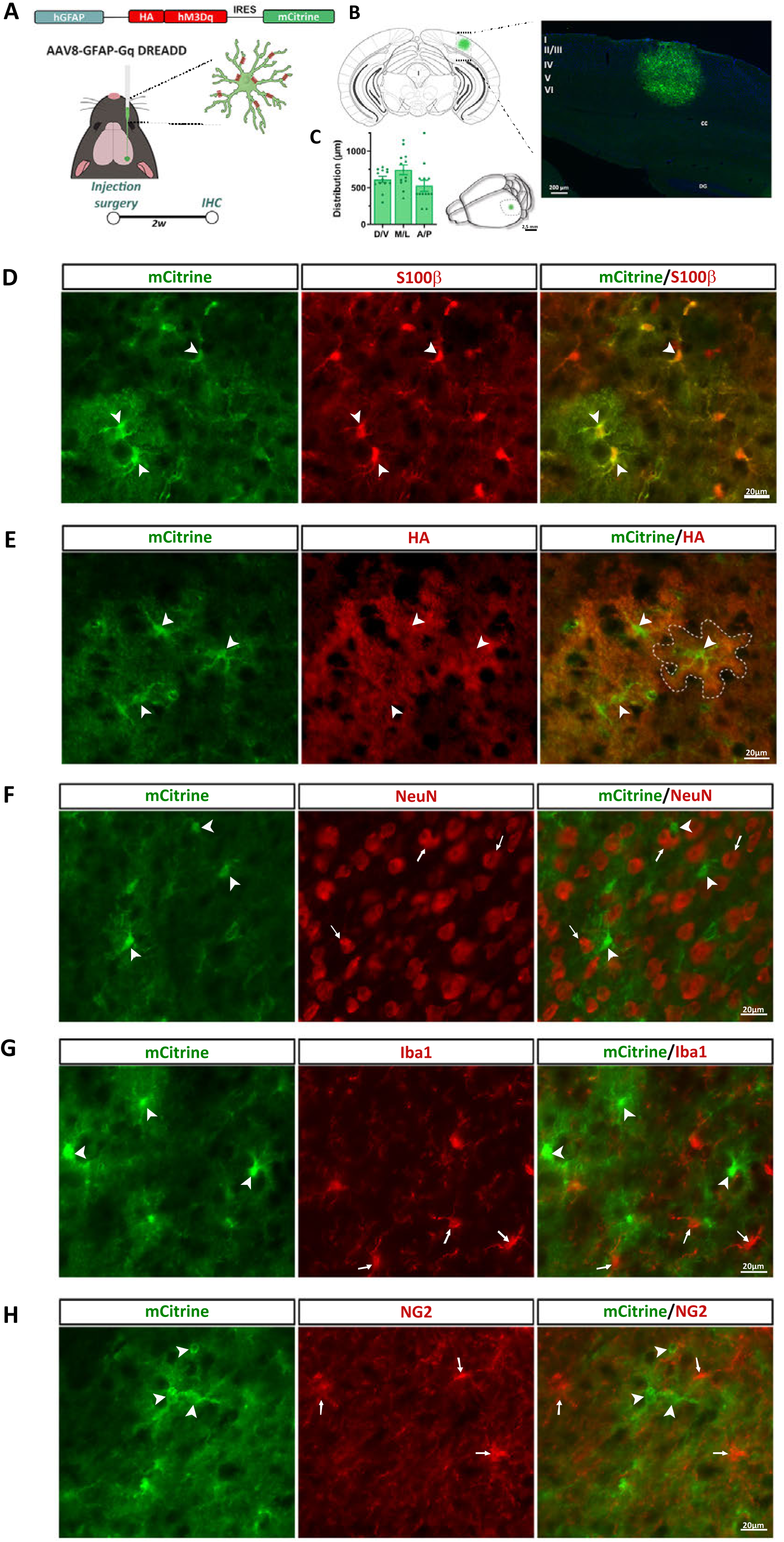
hM3D_q_ receptor is expressed in 73% of visual cortex astrocytes and is specific to V1. **(A)** Molecular construct of the AAV8-GFAP-G_q_ DREADD (top panel); Experimental protocol for cellular characterization: 6-week-old mice were unilaterally injected with AAV8-GFAP-G_q_ DREADD. Tissues are harvested 2 weeks after and IHC is performed. **(B)** mCitrine (green) is expressed in all layers of V1 (Schematic on the left and IHC on the right), Bregma -3,52 mm. **(C)** Quantification, (left graph) and schematic representation (right panel) of mCitrine expression. **(D)** Immunohistochemistry showing that mCitrine (green) is selectively expressed in 73% of astrocytes (S100β, red, arrowheads). **(E)** HA is well expressed in mCitrine^+^ astrocytes, and **(F)** not in neurons (NeuN, red, arrows) or **(G)** microglial cells (Iba1, red, arrows) or **(H)** NG2^+^ cells (NG2, red, arrows). (n=10 mice, n=3 slices/mouse). Abbreviations: cc= corpus callosum; DG= dental gyrus.

To assess whether G_q_ DREADD was functional and induces well the canonical Ca²^+^ signaling in adult V1 astrocytes, we performed two-photon Ca²^+^ imaging in V1 acute slices of the GLAST-CreERT²::GCaMP6f transgenic mouse line. We recently generated and characterized this line and show that the genetically-encoded Ca^2+^ reporter GCaMP6f was specifically expressed in astrocytes of V1 layers^63^. Similarly as above-mentioned experiments, AAV8-GFAP-G_q_DREADD was stereotaxically delivered in V1 of these GLAST-CreERT²::GCaMP6f mice and two week later acute V1 slices were prepared (**Supplementary** Figure 1A,B). CNO (10μM) was bath application reliably triggered intracellular Ca²^+^ elevations in G_q_ DREADD expressing astrocytes, while artificial cerebral spinal fluid (ACSF) did not (**Supplementary** Figure 1C). Taken together, the anatomical and functional results validate the use of AAV8-GFAP-G_q_DREADD for examining the role of chronic astrocytic G_q_ GPCR Ca^2+^ signaling cascades in visual-induced synaptic transmission and LTP in awake mice.

### Chronic astrocytic G_q_ GPCR Ca^2+^ signaling activation depresses LTP magnitude in vivo

To test whether chronic astrocytic G_q_ GPCR Ca^2+^ is sufficient to reshape neuronal networks and impair brain ability to form memory, we used a visual learning paradigm based on experience-dependent LTP (or SRP), which has been reported to exhibit all the features of Hebbian synaptic plasticity^55^ and visual perceptual learning^64^. In this paradigm, VEP long-lasting potentiation was achieved by daily exposure to oriented grating (low level) stimulus^54,64^ (**Figure 2A**). In our experimental design, and prior to visual stimulus exposure, mice were bilaterally injected with the AAV8-GFAP-G_q_ DREADD or artificial cerebrospinal fluid (ACSF for controls) in V1. Two weeks later mice were chronically and daily treated (for 6 weeks) with either CNO (1mg/kg) or saline solution (0.9% NaCl, for controls) (**Figure 2C**). Four groups of mice were thus generated: an experimental group (called DREADD/CNO) and 3 control groups (called DREADD/Saline, ACSF/Saline, and ACSF/CNO). At the end of the chronic treatment period, mice were bilaterally implanted with recording electrodes in V1 layer IV, and 48h later, they were subjected to 1 Hz phase-reversing, full-field, 0.05 cycle/°, 100% contrast, sinusoidal grating stimulus, of 45° orientation during 9 consecutive days (**Figure 2B,C**). On the 10^th^ day, mice were presented with a novel differently oriented stimulus (135°) to confirm that the occurring VEP LTP was dependent on the familiarity with the trained stimulus.

**Figure 2:**
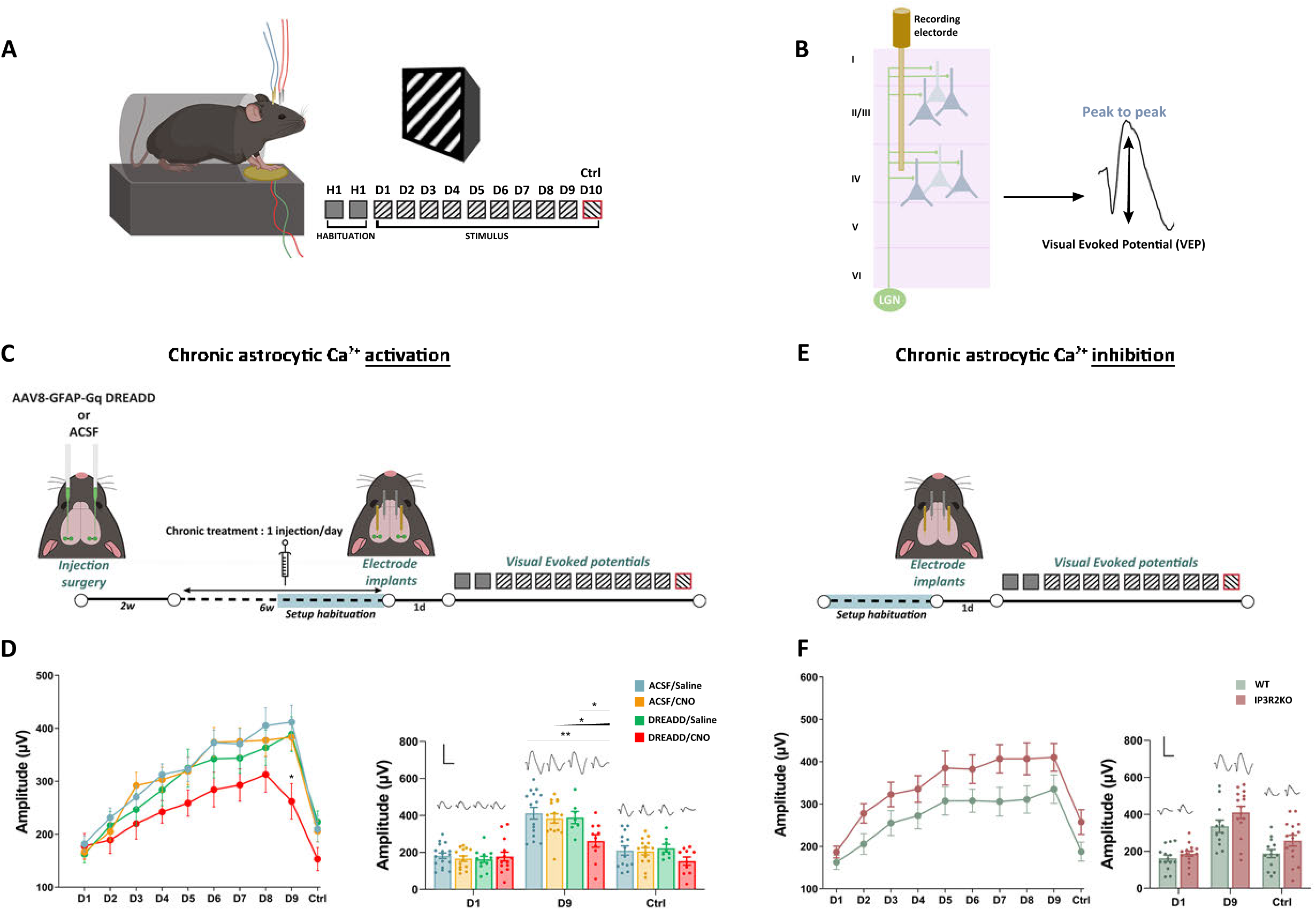
Chronic activation of astrocytic Ca²^+^ signaling impairs SRP formation. **(A)** Experimental design to assess VEP and vidget measurements. Mice are habituated for 2 days to a grey screen, then submitted to the same visual stimulus (45°, black squares) during 9 consecutive days and VEPs and vidgets are recorded. On the 10^th^ day, mice were exposed to novel stimulus with an orientation of 135° (red square). **(B)** Recording electrodes are implanted in V1 layer IV, VEPs are recorded, and peak-to-peak value is extracted. **(C)** Protocol for chronic astrocytic activation: 6-week-old animals are stereotaxically bilaterally injected with the AAV8-GFAP-G_q_ DREADD or ACSF. After 2 weeks, they receive once per day, a chronic 6 week-long treatment either with CNO (1mg/kg) or saline (0.9%). Two days before the end of the treatment, recording electrodes are implanted within V1. 48h later, mice are exposed to the paradigm described in A. Two weeks before the beginning of the VEPs recording, mice are daily handled and habituated to the setup. **(D, F)** Left panel: VEPs amplitude timeline along stimulation days (in µV). Right panel: Average traces showing VEP amplitude on D1 D9 and control D10. n=12-18 electrodes. ANOVA, Two-way repeated measure *p <0.005. **(E)** Protocol for chronic astrocytic blockade: 14-week-old IP_3_R2 KO mice are implanted with recording electrodes and VEPs are measured. All data are mean ± SEM. Abbreviations: ACSF: Artificial Cerebral Spinal Fluid; Ctrl= control; d=day; w=week; WT=Wild type.

On the first day of stimulation, CNO-treated DREADD mice displayed the same VEP amplitude responses as compared to the 3 DREADD/Saline, ACSF/Saline and ACSF/CNO control groups (162.4 ± 16.19µV; 181.8 ± 14.9µV and 166.5 ± 14.62µV respectively; **Supplementary Table 3**). Furthermore, the kinetics of VEP LTP induction in adult mice was confirmed^54^, *i.e.* visual response potentiation was observed at day 2 compared to day 1, and reached a *plateau* at day 6 in the 4 DREADD/Saline, DREADD/CNO, ACSF/Saline and ACSF/CNO groups (**Figure 2D**, **Supplementary Table 3**). Moreover, VEP amplitudes came back to baseline level when mice of the 4 groups were presented with a novel stimulus on day 10. Together, these results indicate that chronic activation of astrocytic G_q_ GPCR signaling is not sufficient to significantly impairs basic visual-evoked excitatory synaptic transmission as well as V1 neuronal network ability to generate LTP (**Figure 2D**). However, even though all 4 mouse groups exhibited VEP potentiation, a tendency to decrease, which becomes significant at day 9 (36 % decrease) in the amplitude of VEP LTP was observed in DREADD/CNO group (VEP amplitude: 312.8 ± 33.6µV) compared to the 3 control groups (ACSF/Saline: 405.1 ± 33.7µV; ACSF/CNO: 377.6 ± 30.1µV; DREADD/Saline: 363.3 ± 33.4µV; mixed effect model, Tukey multiple comparison test, days x treatment interaction p=0.0291; **Figure 2D left and right panels**, **Supplementary Table 3**). This result indicates that chronic activation of astrocytic G_q_ GPCR signaling is sufficient to impair VEP LTP magnitude in a long-lasting manner (*i.e.* days after the end of upregulated astrocytic activation period). Importantly, ACSF/CNO group potentiated similarly compared to the 2 other control groups (ACSF/Saline and DREADD/Saline, **Supplementary Table 3**), showing that CNO application and G_q_ DREADD expression do not trigger non-specific effects in themselves.

Since we observed that chronic upregulated astrocytic G_q_ GPCR signaling induces a decrease in VEP LTP magnitude, we next asked whether astrocytic G_q_ GPCR Ca²^+^ signaling was necessary for this LTP decrease to occur. We reasoned that if chronic astrocytic G_q_ GPCR-mediated Ca²^+^ elevations reshape V1 neuronal network underlying LTP formation, then the chronic removal of astrocytic G_q_ GPCR Ca²^+^ signaling should prevent this neuronal network and LTP impairments. Therefore, the SRP paradigm was performed in IP_3_R2 knockout (KO) mice, enabling the chronic blunting of G_q_ GPCR Ca²^+^ signaling in 100% of astrocytes without affecting neuronal Ca^2+^ responses^65,66^. On day 1, the magnitude of VEPs was similar in IP_3_R2 KO compared to wild type (WT) control mice (WT mice: 162.5 ± 16.8µV; IP_3_R2 KO: 186.9 ± 13.9µV; **Figure 2F**, **Supplementary Table 4**), indicating that dramatic downregulation of global G_q_ GPCR Ca^2+^ signaling in IP_3_R2 KO astrocytes does not affect basic thalamocortical excitatory synaptic transmission. Furthermore, IP_3_R2 KO mice did not show statistically significant alterations of VEP LTP amplitudes compared to WT mice at day 9 [mixed model analysis indicating only a main effect of genotype (p<0.0001); no effect of time (p=0.056) and of interaction between both variables (p=0.40833); **Supplementary Table 4**]. Of note, IP_3_R2 KO mice exhibit a tendency to increase in VEP amplitude across days of stimulation. Together, these results suggest that chronic upregulation of astrocytic G_q_ GPCR Ca^2+^ signaling is not only sufficient (**Figure 2D**, **Supplementary Table 3**) but is also necessary for the decrease in VEP LTP amplitude observed in DREADD/CNO mice.

### Chronic astrocytic G_q_ GPCR Ca²^+^ activation induces microglial cell activation

We assumed that chronic activation of astrocytic G_q_ GPCR Ca^2+^ signaling mimics adult life challenges (*e.g*. chronic stress/inflammation), and thus may lead to astrocyte, but also, microglial cell reactivity since astrocyte-to-microglia communication is now well documented^67,68^. Therefore, we next assessed astrocytic reactivity by measuring GFAP expression level using immunohistochemistry in unilaterally AAV8-GFAP-G_q_DREADD-injected mice treated either with CNO (DREADD/CNO) or saline (DREADD/Saline). (**Figure 3A**). GFAP expression is normally scarce in adult mouse cortex^69^, but can be dramatically enhanced upon injuries or inflammation^70^. Similarly to DREADD/Saline control mice, expression level of GFAP (quantified in DREADD-expressing hemispheres) remained sparse in V1 of DREADD/CNO, indicating that chronic astrocytic G_q_ DREADD activation does not lead to major astrogliosis (**Figure 3B-C**; **Supplementary Table 5**, unpaired t-test; p=0.3860). This result suggest that our experimental protocol rather mimics the early steps of the stress/inflammatory process in the diseased brain when astrocyte gross morphology is not yet altered.

**Figure 3:**
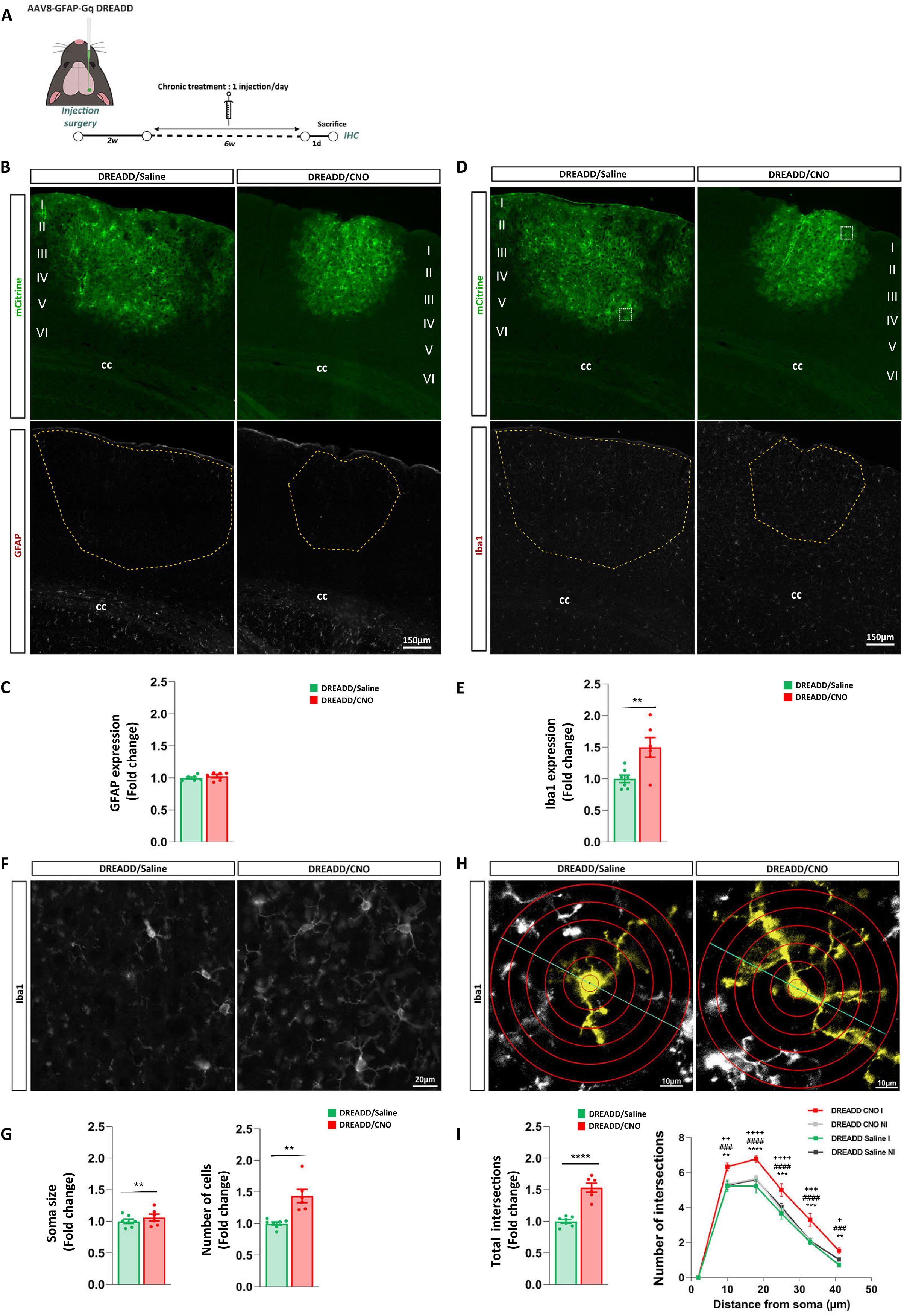
Chronic activation of astrocytic Ca²^+^ signaling triggers microglia activation without astrogliosis. **(A)** Experimental design for immunohistochemistry analysis: 6-week-old animals are stereotaxically unilaterally injected with the AAV8-GFAP-G_q_ DREADD. After 2 weeks, they receive once per day, a chronic 6 week-long treatment either with CNO (1mg/kg) or saline (0,9%). 1 day after the end of the treatment, mice are sacrificed, and brain harvested for IHC. **(B, D)** Representative epifluorescence images of DREADD/Saline (left) and DREADD/CNO (right) immunohistochemistry in V1 stained for mCitrine (green, top panels) and for GFAP **(B**, bottom panel, white) and Iba1 **(D,** bottom panel, white) (). Yellow dashed lines represent the analyzed region where the DREADD is expressed. **(C-E)** GFAP **(C)** and Iba1 **(E)** expression level fold change within V1 ROI. **(F)** Representative images of Iba1^+^ microglial cell density in DREADD/Saline (left) and DREADD/CNO mice (right). **(G)** Soma size measurement of Iba1^+^ cells (left panel) and number of Iba1^+^ cells (right graph) in DREADD/Saline compared to DREADD/CNO mice. **(H)** Representative epifluorescence images showing Sholl analysis on microglia (yellow) from DREADD/Saline (left) and DREADD/CNO mice (right). **(I)** Quantification of the total number of microglial intersections (fold change) in DREADD/Saline and DREADD/CNO mice (left graph) (data normalized to NI hemisphere); and Sholl analysis plot (right graph) showing the number of intersections of microglial branches with each circle depending on the distance from the soma. Data were measured for each injected hemisphere in DREADD/Saline and DREADD/CNO mice and their corresponding non-injected control hemisphere. Data were tested for statistical differences between groups using two-ways ANOVA and Bonferroni post-hoc testing, or unpaired, non-parametric Mann-Whitney test when only two groups. (* = p<0.05, ** = p<0.01 between CNO I and CNO NI; #= p<0.05, ## = p<0.01, ### = p<0.001, #### = p<0.0001 between CNO I and Saline I; ^+^ = p<0.05, ^++^ = p<0.01, ^+++^ = p<0.001, ^++++^ = p<0.0001 between CNO I and Saline NI. All data are mean ± SEM. Abbreviations: cc = corpus callosum, I=injected hemisphere; NI= non-injected hemisphere. I II III IV V VI represent V1 cortical layers.

Microglial reactivity is also a hallmark of inflammation, and is associated with cell proliferation, changes in soma size and morphology^71^. To further assess the influence of chronic astrocytic G_q_ GPCR activation in microglia function, the expression level of Iba1, a gold standard marker of microglia reactivity, was measured. A statistically significant 50% increase in the overall Iba1 expression level was observed in DREADD/CNO compared to DREADD/Saline mice (**Figure 3D-E**, **Supplementary Table 5**, unpaired t-test; p=0.0086). Such increase was due to both a 43% increase in the number of microglial cells (**Figure 3F-G**, **Supplementary Table 5**, unpaired t-test, p= 0.0011) and a 16% enlarged soma size (**Figure 3F-G**, **Supplementary Table 5**, unpaired t-test; p=0.0042) in DREADD/CNO group compared to the control group. Because, chronic stress^72^, inflammation^73^ as well as early form of dementia^74^, neurodegenerative diseases^75^ or β-amyloid plaques^76^ have been showed to trigger (or be associated with) microglia hyper-ramification, we evaluated microglial morphology. Sholl analysis demonstrated an overall 53% increase in the cumulative total process intersection number in DREADD/CNO mice compared to DREADD/Saline controls (**Figure 3H,I**, **Supplementary Table 5**, unpaired t-test; p<0.0001). Furthermore, the number of process intersections was further investigated in the AAV8-GFAP-G_q_DREADD-injected hemispheres (called DREADD I) *versus* non-injected internal control hemispheres (called DREADD NI) from DREADD/CNO and DREADD/saline mice. A high increase (of 60 to 70%) in the number of intersections was observed in AAV-injected and CNO-treated V1 (DREADD/CNO I) compared to the 3 control groups [AAV-injected and saline-treated V1 (DREADD/Saline I), non-injected and CNO-treated V1 (DREADD/CNO NI), and non-injected and saline-treated V1 (DREADD/saline NI) (**Figure 3I right panel, Supplementary Table 5**, two-way repeated measure ANOVA followed by Tukey’s multiple comparison test; distance x number of intersections, p= 0.0197). The fact that no difference in process intersection number was detectable between the 3 controls groups (DREADD/Saline I, DREADD/CNO NI and DREADD/saline NI) demonstrated that chronic (6 week-long) daily CNO (1mg/kg) administration does not lead to off-target effects and confounding results (**Figure 3I right panel, Supplementary Table 5**,). Therefore, our data show that chronic astrocytic G_q_ GPCR Ca^2+^ signaling leads to microglial cells, the main brain innate immune cells, to display increased reactivity toward a non-macrophage-like phenotype. This suggest the occurrence of a G_q_ GPCR-mediated astrocyte-to-microglia communication, which could mimic the early sequences of inflammatory events contributing to synapse loss in neurodegenerative diseases, including AD.

### Chronic astrocytic G_q_ GPCR Ca²^+^ activation leads to alterations of interleukin-33 expression levels

To test whether chronic upregulation of astrocytic G_q_ GPCR Ca^2+^ signaling could alter the secretion of some astrocyte-specific factors known to induce microglia hyper-ramification, we quantified interleukin-33 (IL-33) expression level in the same mouse groups as described in Figure 3. The selection of IL-33 as a candidate molecule was based on the facts that (i) IL-33 is a cellular alarmin and is released from nuclear stores of producing cells after an inflammatory process^77^, (ii) IL-33 is mainly expressed in CNS astrocytes^56,78,79^, (iii) during brain development, astrocytic IL-33 is secreted to act on its interleukin 1 receptor-like 1 (ST2), which is selectively expressed in microglia plasma membrane^56^, (iv) hippocampal IL-33 infusion triggers increased microglial density and hyper-ramification, associated with upregulated IL-1β production and an alteration in long-term memory formation^80^, and (v) genetic polymorphisms within IL-33 gene have been associated with AD risk^57^. We first determined whether V1 astrocytes express nuclear IL-33; we detected this interleukin in 57 % of the astrocyte population within all V1 layers, with an enrichment (63 %) in deep layers IV-VI astrocytes (n= 2 WT adult mice).

To next determine whether IL-33 secretion was upregulated upon chronic astrocytic G_q_ GPCR activation, the number of V1 astrocytes expressing nuclear IL-33 was quantified in DREADD/CNO *versus* DREADD/Saline mice. No change, neither in overall IL-33 expression level nor in IL-33-expressing cell number, was found within V1 layers I-V between both mouse groups (**Figure 4A,B** yellow-dashed-line region quantified in **4B** left, **Supplementary Table 6**, Two-tailed Mann Whitney, p = 0.1807 and middle panel: Unpaired t-test, p=0.7368). However, we noticed a downward trend in IL-33 expression level in deeper V1 layers IV and V of DREADD/CNO mice compared to DREADD/Saline controls (**Figure 4C,D**, **Supplementary Table 6**, Two-way ANOVA, Distance (μm) x Treatment, p=0.6709 interaction, p<0.0001, Bonferroni’s multiple comparisons post-hoc test). This observation led us to quantify, only in V1 layer IV, the number of cells expressing IL-33 in their nucleus. A 23% decrease in such number was detected (**Figure 4A** blue-dashed line region, **Figure 4E**, **Supplementary Table 6**, Unpaired t-test, p<0.0001), suggesting that IL-33 release from astrocytic nucleus into the extracellular space^81^ is upregulated upon chronic astrocytic G_q_ GPCR activation, which might induce microglial reactivity and hyper-ramification (**Figure 3**).

**Figure 4:**
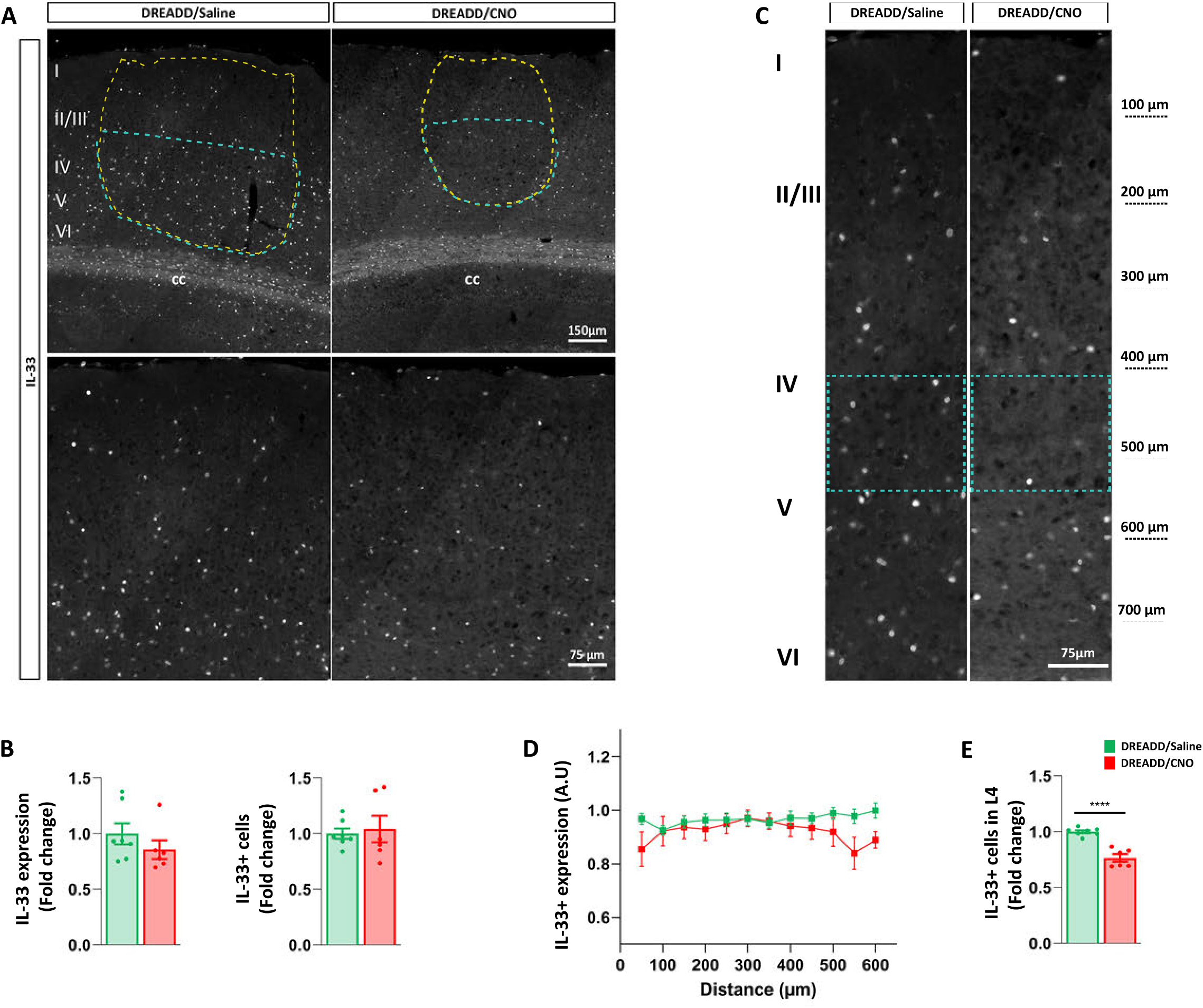
Chronic activation of astrocytic Ca²^+^ signaling leads to IL-33 modulation. **(A)** Representative epifluorescence images of DREADD/Saline (left) and DREADD/CNO (right) immunohistochemistry in V1 stained for IL-33 (top panels, full field and magnification of layer I, II/III on bottom panels). **(B)** left panel graph: quantification of IL-33 expression (mean grey value, fold change) within the entire ROI (represented by the yellow dashed lines in A); right panel graph: quantification of the number of IL33^+^ cells (fold change) within the entire ROI (represented by the yellow dashed lines in A). **(C)** Representative epifluorescence images of V1 cortical layers from DREADD/Saline (left) and DREADD/CNO (right) stained for IL-33. **(D)** Quantification of plot line-scan mean density measurements for IL-33-immunostaining across the entire V1 (normalized to non-injected hemisphere values). Blue dashed rectangles represent V1 layer IV. **(E)** Quantification of the number of IL33^+^ cells (fold change) only from layer IV to the end of the viral spread. All data are mean ± SEM.. An unpaired, non-parametric Mann-Whitney test has been performed. *p<0.05, **p<0.01, ***p<0.005, **** = p<0.0001. Abbreviations: I II III IV V VI represent V1 cortical layers. n=6 CNO-treated DREADD mice; n=7 Saline-treated DREADD mice.

### Chronic astrocytic G_q_ GPCR Ca²^+^ activation does not modulate the number of excitatory thalamocortical synapses within V1 Layer IV

Given that chronic astrocytic G_q_ GPCR Ca^2+^ activation led to a decrease in excitatory thalamocortical synaptic LTP magnitude (**Figure 2**), as well as microglial reactivity (**Figure 3**) possibly mediated through upregulated IL-33 release from layer IV astrocytes (**Figure 4**), we hypothesized that LTP impairment was due to an excess of microglial thalamocortical synapse elimination in V1 layer IV. Indeed, in the developing thalamus and dLGN, astrocytic IL-33 release has been found to trigger microglial-mediated synapse pruning^56^. It is thus reasonable to think that such phenomenon could be reactivated in adult V1 upon chronic upregulation of astrocytic G_q_ GPCR Ca^2+^ signaling. To investigate this hypothesis, we used the same mouse groups as previously described (**Figures 3 and 4**), to assess the number of excitatory thalamocortical synapses, which selectively express the vesicular glutamate transporter 2 (VGlut2) marker in their presynaptic compartments (**Figure 5A**, left scheme). Using Western blot, no statistically significant changes in VGlut2 expression levels was observed between DREADD/CNO and DREADD/Saline mice (**Figure 5A-C**, **Supplementary table 7**, unpaired t-test, p=0.0866), suggesting that the number of V1 excitatory thalamocortical synapses was not affected by chronic astrocytic activation. In agreement, expression levels of the postsynaptic density protein 95 (PSD-95), a marker of excitatory post-synaptic compartments, was unchanged in V1 of DREADD/CNO compared to DREADD/saline mice (**Figure 5C**, **Supplementary table 7**, unpaired t-test, p=0.0866). To further evaluate the number of excitatory thalamocortical synapse restricted to V1 layer IV, we used immunohistochemistry to detect VGluT2-expressing thalamocortical presynaptic compartments and Homer-1-expressing postsynaptic excitatory terminals (**Figure 5D-F**). High resolution confocal microscopy did not show any changes in neither the number of both VGlut2-(Mann Whithney test, p=0.6278), and Homer1-expressing puncta (unpaired t-test p=0,7187), nor in the juxtaposition of these puncta (Unpaired t-testp=0.6325) in DREADD/CNO compared to DREADD/Saline mice (**Figure 5F**, **Supplementary table 7**), demonstrating that chronic astrocytic G_q_ GPCR Ca²^+^ activation does not alter the number of excitatory thalamocortical synapses within V1 layer IV. Therefore, the astrocytic G_q_ GPCR-evoked reduction in LTP magnitude (**Figure 2**) is likely due to another mechanism, currently under investigation than to microglia-mediated removal of layer IV excitatory synapses.

**Figure 5:**
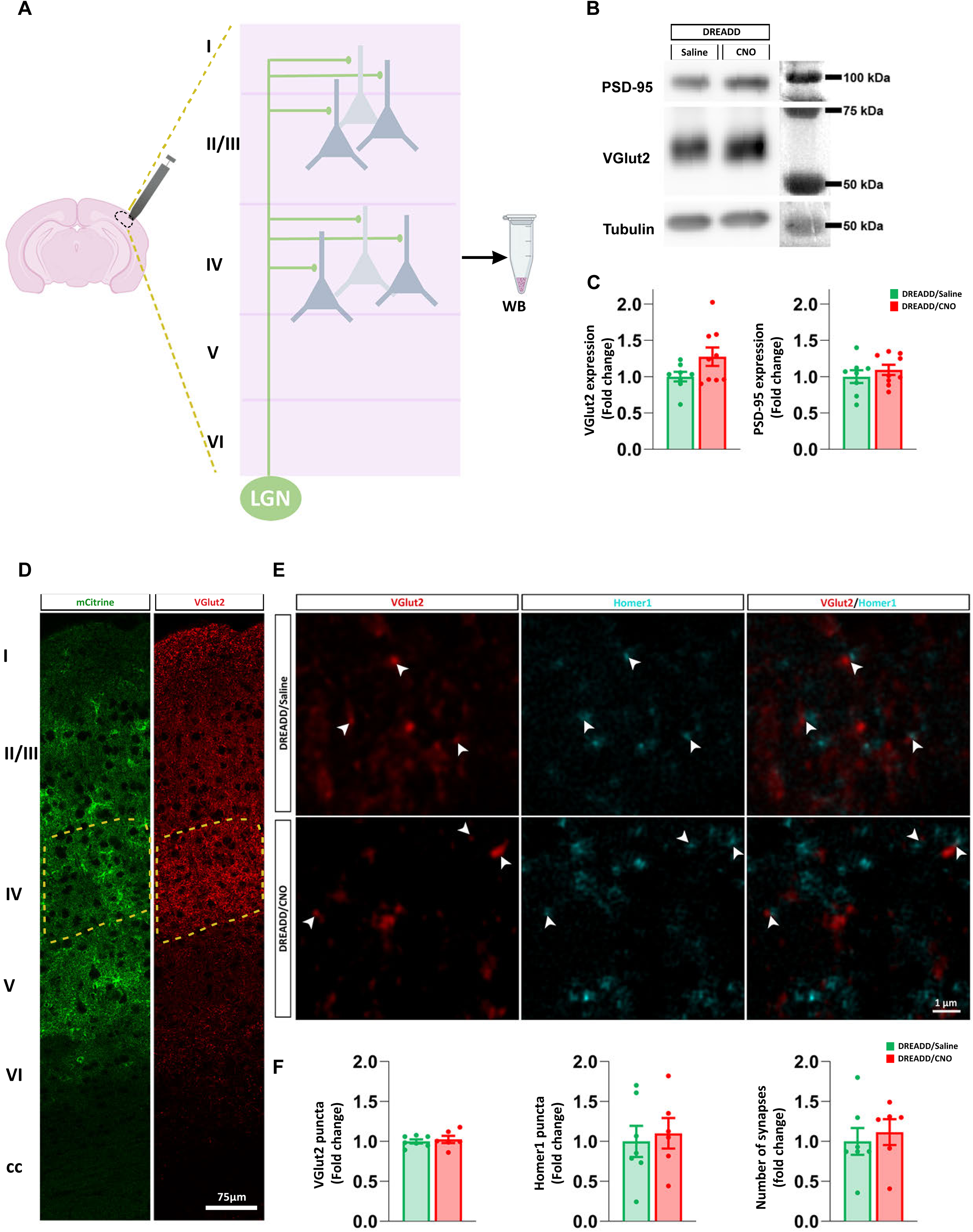
Chronic activation of astrocytic Ca²^+^ signaling does not affect excitatory synapse number. **(A)** Representative schematic of a mouse brain where 2mm of all V1 layers are “punched” to perform western blots. LGN afferences (green) onto layer IV and layer I pyramidal neurons represent VGlut2^+^ inputs. **(B)** WB representative images of PSD-95 and VGlut2 expression in V1 from DREADD/Saline and DREADD/CNO. Tubulin: loading control. **(C)** WB quantifications of VGlut2 expression (left graph, fold change) and PSD-95 (right graph, fold change) from DREADD/Saline and DREADD/CNO mice. **(D)** Confocal images showing mCitrine (left image), VGlut2 (right image) through V1 cortical layers. Increased VGlut2 staining shows thalamocortical main excitatory projections in V1 layer I and IV (dashed yellow rectangle). **(E)** Representative confocal images taken from layer IV mCitrine-expressing V1 region (defined by the dashed yellow rectangle from **D**), showing excitatory VGlut2-immunoreactive puncta (red, left panels), Homer1-immunoreactive puncta (blue, middle panels) and merged staining (right panels) from DREADD/Saline and DREADD/CNO mice. White arrowheads indicate detectable excitatory thalamocortical synapses, formed by both VGlut2^+^ pre-and Homer1^+^ post-synaptic compartments. **(F)** Quantification of VGlut2-and Homer1 -immunoreactive puncta, and synapses formed by both VGlut2 -and Homer 1-immunoreactive puncta (fold change). Data are shown as mean ±SEM. An unpaired, non-parametric Mann-Whitney test has been performed.

## Discussion

In this study, we have investigated the role of astrocytic G_q_ GPCR Ca^2+^ signaling in synaptic transmission and plasticity within the adult mouse V1. We found that chronic upregulation of this Ca^2+^ signaling pathway is sufficient to reduce by ∼ 36% the formation of thalamocortical LTP (SRP) in V1 layer IV. Chronic abolishment of the main G_q_ GPCR Ca^2+^ elevations using the IP_3_R2 KO mouse model provided further evidence to support the idea that sustained GPCR-mediated Ca^2+^ elevations are not only sufficient but might also be necessary to alter LTP formation. Moreover, astrocytic G_q_ GPCR Ca^2+^ signaling evoked a dramatic increase in microglia reactive state, including a ∼ 40-50% enhancement in microglial Iba1 expression level, cell number, and process intersection number, which were associated with a ∼ 23 % decrease in IL-33 nuclear expression in layer IV astrocytes.

These results have profound implications for AD, as upregulated astrocytic P2Y_1_R and mGluR5-mediated Ca^2+^ signaling have recently been reported to play an important role in the onset or development of AD^30–34^. However, whether such abnormally increased astrocytic G_q_ GPCR Ca^2+^ elevations are a consequence (or a cause) of AD-associated LTP and memory deficits is unknown. Our study, showing that chronically increased astrocytic G_q_ GPCR Ca^2+^ signaling induces alterations of visually-evoked LTP formation, provides a first a piece of evidence that upregulation of astrocytic G_q_ GPCR Ca^2+^, in itself, can be a cause (rather than a consequence) of AD-like symptom onset. Therefore, our results support our original hypothesis that during adult life challenges, chronic stress/inflammation-induced upregulation of astrocytic G_q_ GPCR Ca^2+^, could contribute to the early pathogenic cascades of events underlying neurodegeneration in AD.

However, our experimental protocol, aiming at mimicking such chronic stress/inflammation-induced upregulation of astrocytic G_q_ GPCR Ca^2+^, did not lead to obvious astrogliosis. This is in agreement with a recently posted article^82^, showing that chronic (7 week-long) G_q_ DREADD activation in mouse cortical and hippocampal astrocytes, does not alter the expression level of genes encoding GFAP and neurotoxic A1 astrocyte proteins. Interestingly, other astrocytic genes encoding proteins involved in GPCR signaling and Ca^2+^ homeostasis were upregulated, with no major transcriptional changes observed in the neurons^82^.

Furthermore, our study has shown that chronic overactivation of astrocytic G_q_ GPCR Ca^2+^ signaling triggered microglial reactivity, hyper-ramification and proliferation, suggesting that physiological homeostatic mechanisms are lost and pro-inflammatory responses are upregulated^75^. Our data are the first to show that astrocytic G_q_ GPCR Ca^2+^ signaling is a key player in astrocyte-to-microglia communication, which has important implication in brain inflammation and diseases. Interestingly, hyper-ramified microglia phenotype is typically found in chronic diseases, such as multiple sclerosis (MS)^73^, PTSD^83^, chronic stress, inflammation^80^, anxiety or depression^72,84,85^, AD^76^ or PD^75^ and other NDs. Such “stress-primed” hyper-ramified morphology may increase the frequency of microglia-synaptic contacts in V1^86^, and thus can potentially alter visually-evoked LTP (SRP) formation. The first obvious, and well documented way for microglia to alter LTP is *via* enhanced phagocytosis of excitatory synapses^56^. The fact that the number of thalamocortical excitatory synapses was not changed upon chronic upregulated astrocytic G_q_ GPCR Ca^2+^ signaling (**Figure 5D-F**), suggest that other synapses (*e.g.* inhibitory synapses^87^ rather than excitatory synapses) may be targeted by microglia.

The chronic activation of astrocytic G_q_ GPCR Ca^2+^ signaling led to decreased number of V1 layer IV astrocytes expressing IL-33 in their nucleus, suggesting an increased IL-33 release into the extracellular space^81^. Such decrease is associated with microglial activation, proliferation and hyper-ramified state^80^, which is in agreement with the idea that astrocytic IL-33 is a key astrocytic mediator resulting in microglial reactive morphology not only in the developing brain^56^, but also in the adult brain^80^. Because studies have showed that (i) *IL-33* polymorphisms were associated with AD risk^57^ and that (ii) increase in both IL-33 or its receptor ST2 expression levels are found around astrocytes and Aβ amyloid plaques in AD patients^58^, our data suggest the possibility that chronic and upregulated astrocytic G_q_ GPCR Ca^2+^ signaling contributes to these aspects of AD pathogenesis.

Finally, and importantly, the altered astrocytic G_q_ GCPR-mediated LTP (SRP) formation here reported, may potentially be involved in the visual perception and object recognition deficits reported in AD patients^42,88,89^. During the course of our research, a study^90^ has shown that the amplitude of such LTP (SRP) was decreased in the rTg4510 model of tauopathy, thus phenocopying our data and strongly supporting such hypothesis. Therefore, impairments in this visual LTP (reflecting a form of visual perceptual memory exclusively stored in V1^64^, could represent a potential functional marker for predicting the development of dementia, and thus the severity and progression of cognitive and the early sensory symptoms associated with AD^41–51^, but also other neurodegenerative disease^91–93^. Furthermore, our date might help understanding better whether and how astrocytic GPCR signaling is involved in the physiopathology underlying LTP (SRP).

In conclusion, and to the best of our knowledge, our findings are the first to demonstrate potential roles for astrocytic G_q_ GPCR signaling in V1 LTP formation and microglial activity state, possibly through an IL-33 pathway. Although caution should be exercised in extrapolation of our findings to other systems and human physiology or pathology (*e.g.* AD), they provide evidence for an active role of astrocytic G_q_ GPCR *in vivo*. GPCRs are a principal target of many therapeutic agents, and it is conceivable that astrocytic GPCRs represent interesting therapeutic candidates for treating the diseased brain. Thus a better understanding of how alterations in astrocyte G_q_ GPCR function disturb surrounding neuronal and microglial cells *in vivo* would be crucial to understand normal physiology and disease evolution.

## Author contributions

EM and CA designed the experiments. EM performed electrophysiology, Ca^2+^ imaging, and immunohistochemistry experiments, analyzed and graphed the results. SA performed stereotaxic injections for immunohistochemistry and Western blot analysis. BR and SG performed Western blot experiments, analyzed and graphed the data. SA and SG took care of the mouse colony. JS helped and advised for *in vivo* electrophysiological recording and data analysis. CA conceived, administrated and supervised the project. EM and CA wrote the manuscript. All authors read and commented the article.

## Acknowledgments

We gratefully acknowledge S. Picaud, A. Bessis, C. Levenes and C. Meunier for valuable discussions and feedback; J.M. Andrieu for sharing laboratory and office spaces; F. Charbonnier for sharing pieces of equipment; and the BioMedTech mouse core facility for hosting our mouse colony.

## Supplementary Information

**Supplementary Figure 1:**
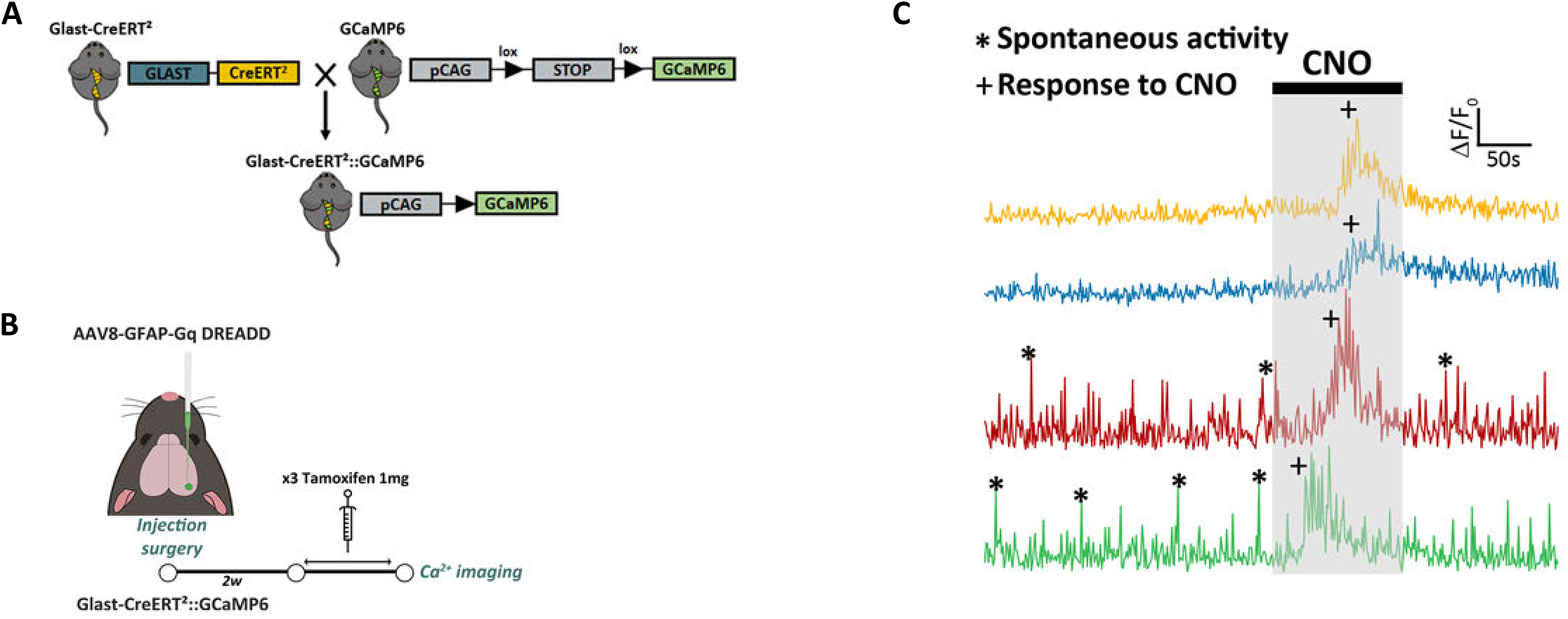
hM3D_q_ is functional in astrocytes. **(A)** Illustration of the different steps to obtain the GLAST-Cre ERT²::GCaMP6 mouse expressing the GCaMP6, (a green calcium indicator) specifically in astrocytes. **(B)** Protocol used for the functional characterization of the G_q_ DREADD. **(C)** 2-photon calcium imaging shows spontaneous and CNO-induced Ca^2+^ increases in DREADD-expressing astrocytes (n=4 cells).

**Supplementary Figure 2:**
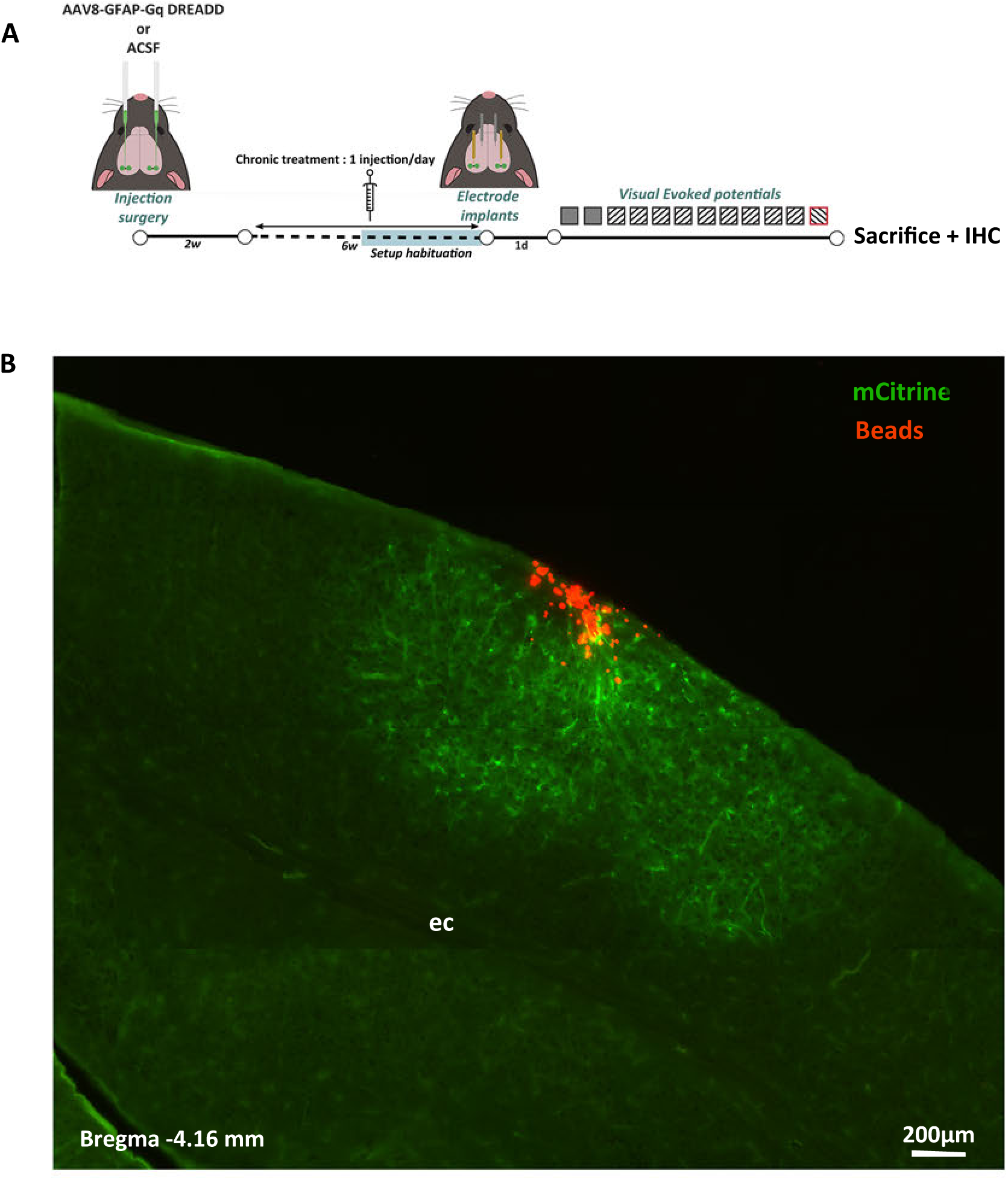
Recording electrodes are dipped in micro-fluorescent beads to check electrode emplacement. **(A)** Experimental protocol: during implantation surgery, recording electrodes are dipped in fluorescent beads. At the end of VEPs recordings, mice are sacrificed, and electrode placement is checked for all mice. **(B)** Representative epifluorescence images showing that electrode track (red beads) is in the middle of mCitrine^+^ astrocytes (green). Abbreviations: ec= external capsule

**Supplementary Table 1:** List of antibodies.

| Primary antibodies | Species | Reference | Company | Dilution |
| --- | --- | --- | --- | --- |
| Anti-GFP | Chicken | A10262 | Invitrogen | 1:1000 |
| Anti-Iba1 | Rabbit | 019-19741 | Wako | 1:1000 |
| Anti-GFAP | Mouse | MAB360 | Millipore | 1:1000 |
| Anti S100b | Rabbit | Ab52642 | Abcam | 1 :1000 |
| Anti-NeuN | Guinea Pig | ABN90 | Millipore | 1 :1000 |
| Anti-HA | Rabbit | 37245 | Cell signaling | 1 :1000 |
| Anti-IL-33 | Goat | AF3626 | R&D | 1 :500 |
| Anti Homer1 | Rabbit | 160003 | Synaptic System | 1 :300 |
| Anti VGlut2 | Guinea Pig | AB2251-I | Millipore | 1 :1000 (IHC)<br>1 :10000 (WB) |
| Anti-PSD-95 | Rabbit | 51-6900 | Invitrogen | 1 :1000 |
| Anti-tubulin | Mouse | T6074-200UL | Sigma | 1 :20000 |
| Secondary Antibodies (fluorescent) | Species | Reference | Company | Dilution |
| Alexa Fluor® 488 anti-Chicken | Goat | A-11039 | Invitrogen | 1:1000 |
| Alexa Fluor® 546 anti-Rabbit | Goat | A-11035 | Invitrogen | 1:1000 |
| Alexa Fluor® 568 anti-Mouse | Goat | A-11030 | Invitrogen | 1:1000 |
| Alexa Fluor® 546 Guinea Pig | Goat | A-11074 | Invitrogen | 1:1000 |
| Alexa Fluor® 546 donkey anti-Goat | Donkey | A-11056 | Invitrogen | 1:1000 |
| Brilliant Violet 421 donkey anti-rabbit | Donkey | AB_2651108 | Jackson Immunoresearch | 1 :500 |
| Secondary Antibodies (HRP) | Species | Reference | Company | Dilution |
| HRP Goat anti-Rabbit IgG | Goat | 111-035-003 | Jackson Immunoresearch | 1:10000 |
| HRP Goat Anti-Guinea Pig | Goat | 106-035-003 | Jackson Immunoresearch | 1:10000 |
| HRP Goat Anti-Mouse | Goat | 115-035-003 | Jackson Immunoresearch | 1:10000 |

## Detailed data and statistics

**Supplementary Table 2:**
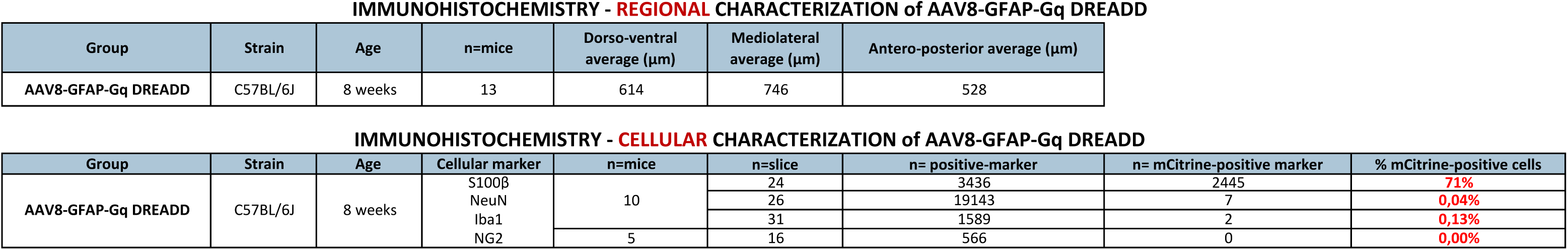
Detailed data and statistics: Regional and functional characterization of AAV8-GFAP-G_q_ DREADD. (Related to Figure 1)

**Supplementary Table 3:**
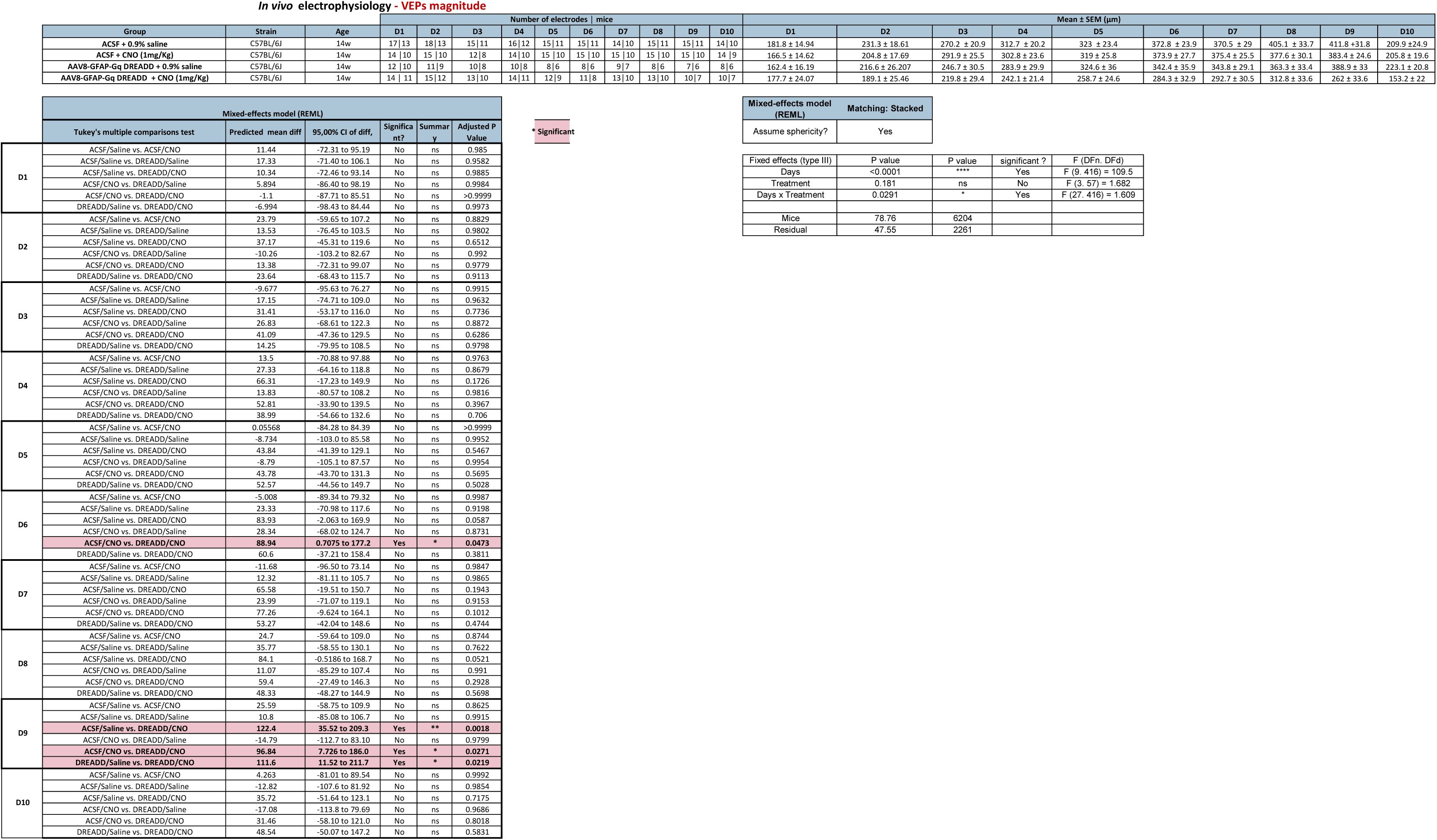
Detailed data and statistics: *in vivo* electrophysiology - VEPs-LTP magnitude of ACSF/Saline, ACSF/CNO, DREADD/Saline and DREADD/CNO mouse groups. (Related to Figure 2)

**Supplementary Table 4:**
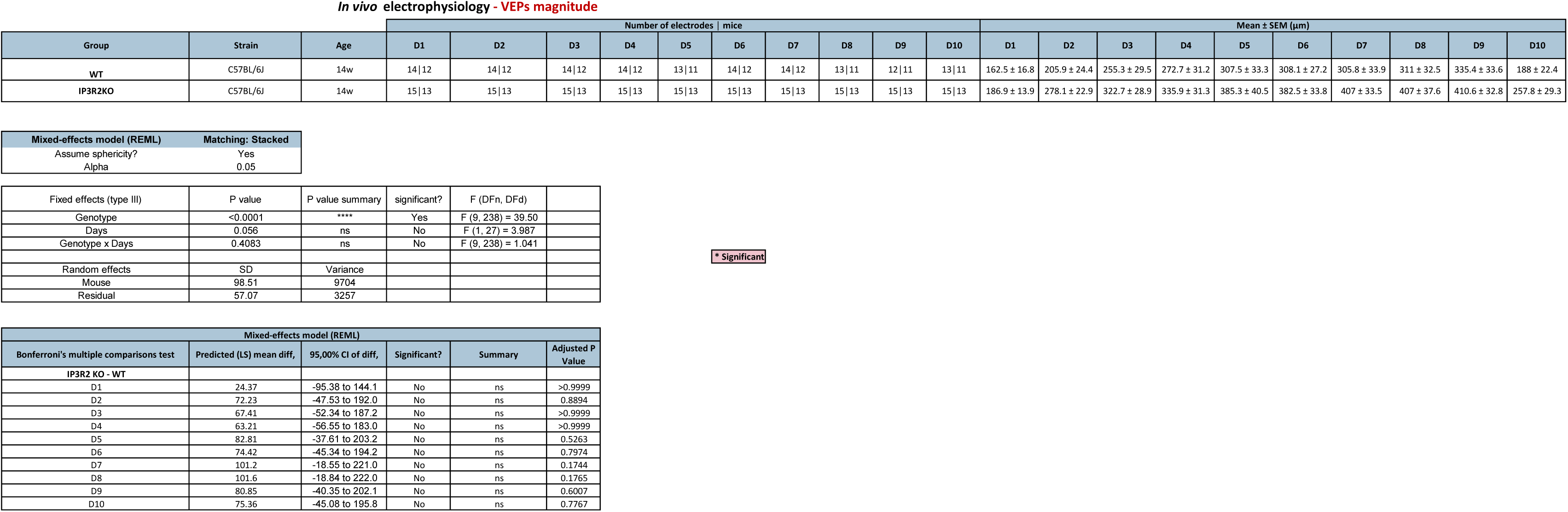
Detailed data and statistics: *in vivo* electrophysiology VEPs-LTP magnitude of WT and IP_3_R2 KO mouse groups. (Related to Figure 2)

**Supplementary Table 5:**
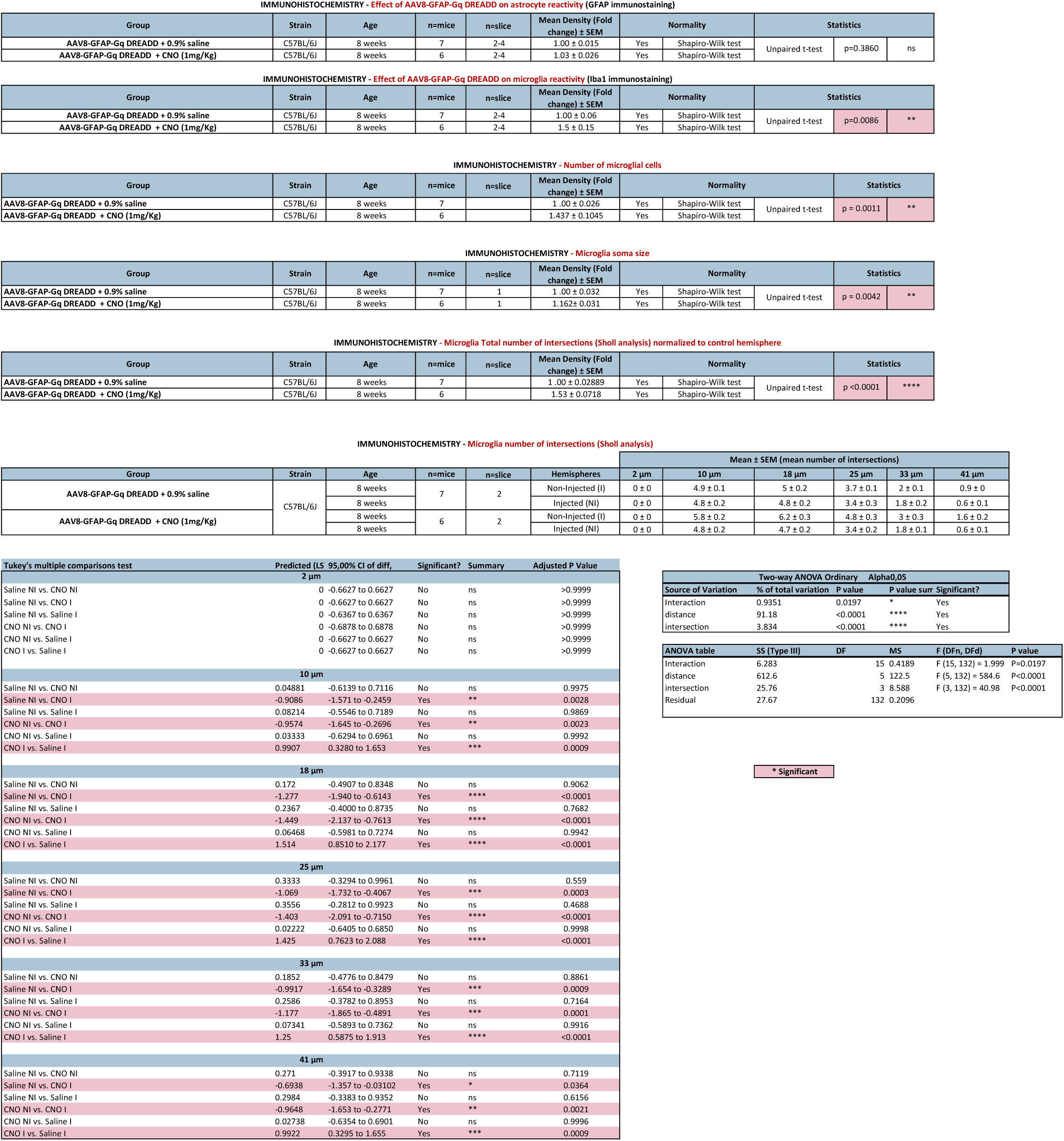
Detailed data and statistics: GFAP and Iba1 quantification in DREADD/Saline and DREADD/CNO mouse groups. (Related to Figure 3)

**Supplementary Table 6:**
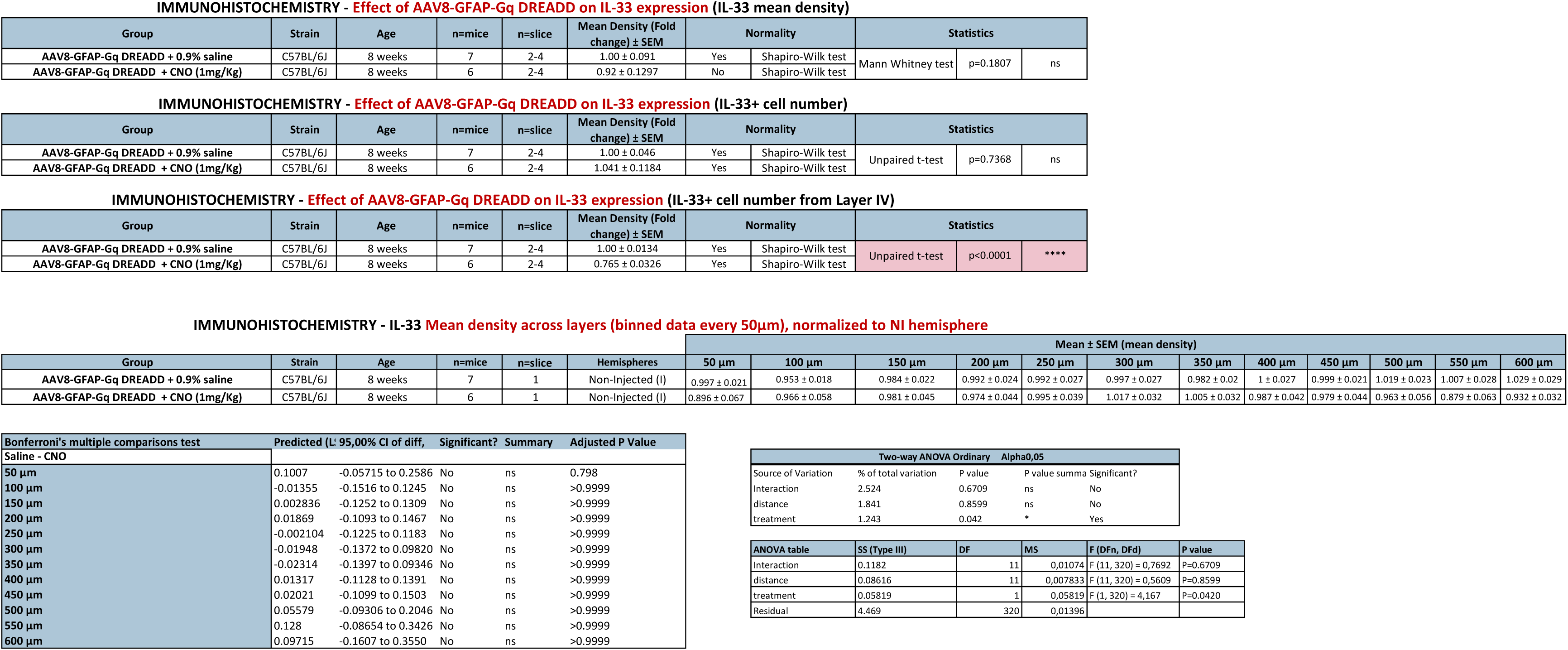
Detailed data and statistics: IL-33 quantification in DREADD/Saline and DREADD/CNO mouse groups. (Related to Figure 4)

**Supplementary Table 7:**
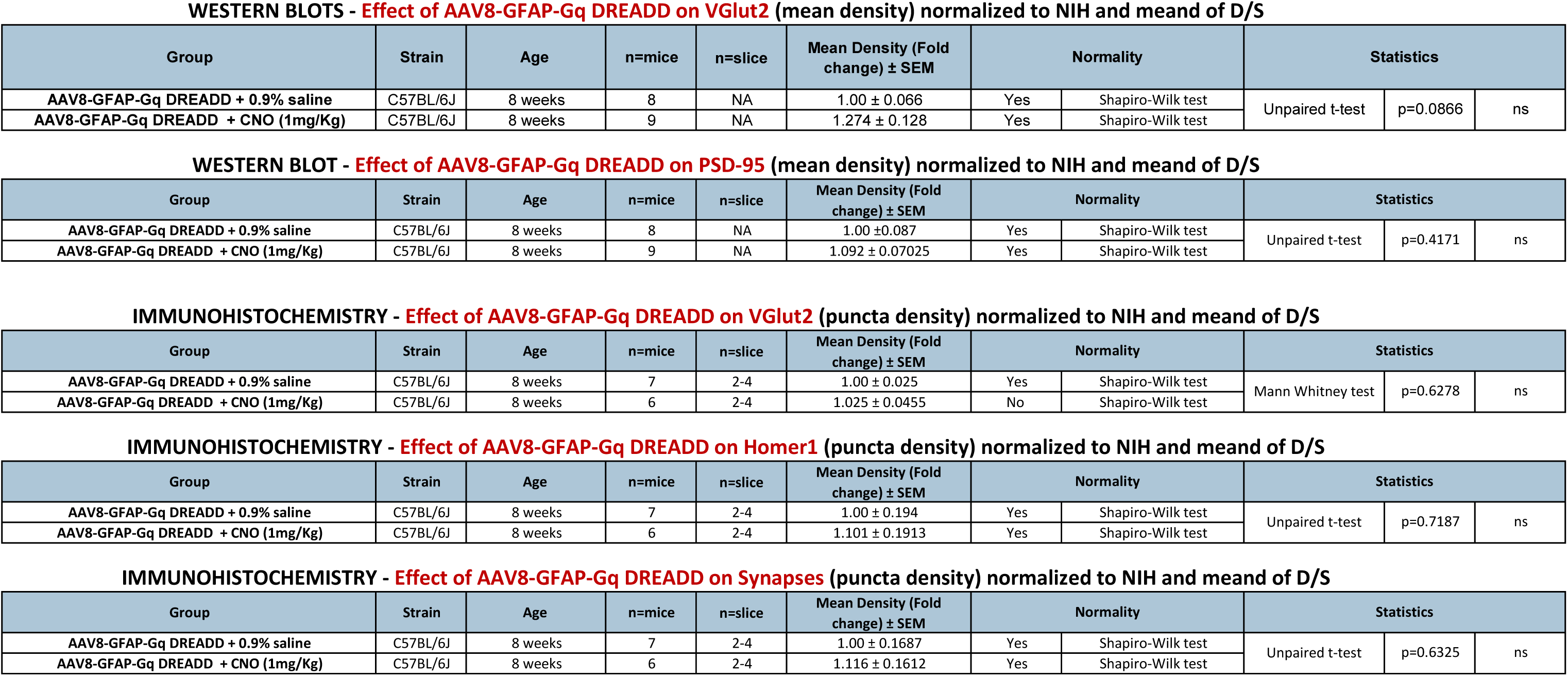
Detailed data and statistics: Synapse quantification in DREADD/Saline and DREADD/CNO mouse groups. (Related to Figure 5)

